# Fission of megamitochondria into multiple smaller well-defined mitochondria in the ageing zebrafish retina

**DOI:** 10.1101/2021.12.09.471958

**Authors:** Thomas Burgoyne, Maria Toms, Chris Way, Dhani Tracey-White, Clare E. Futter, Mariya Moosajee

**Affiliations:** UCL Institute of Ophthalmology, University College London, London EC1V 9EL, UK; Royal Brompton Hospital, Guy’s and St Thomas’ NHS Foundation Trust, London SW3 6NP, UK; The Francis Crick Institute, 1 Midland Road, London NW1 1AT, UK; Department of Ophthalmology, Great Ormond Street Hospital for Children NHS Foundation Trust, London WC1N 3JH, UK; Department of Genetics, Moorfields Eye Hospital NHS Foundation Trust, London EC1V 2PD, UK

**Author notes:** Joint first author (Thomas Burgoyne, Maria Toms). **Corresponding author**, Professor Mariya Moosajee, Development, Ageing and Disease, UCL Institute of Ophthalmology, 11-43 Bath Street, London, EC1V 9EL, United Kingdom.

## Abstract

Mitochondria are essential adenosine triphosphate (ATP)-generating cellular organelles. In the retina, they are highly numerous in the photoreceptors and retinal pigment epithelium (RPE) due to their high energetic requirements. Fission and fusion of the mitochondria within these cells allow them to adapt to changing demands over the lifespan of the organism. Using transmission electron microscopy, we examined the mitochondrial ultrastructure of zebrafish photoreceptors and RPE from 5 days post fertilisation (dpf) through to late adulthood (3 years). Notably, mitochondria in the youngest animals were large and irregular shaped with a loose cristae architecture, but by 8 dpf they had reduced in size and expanded in number with more defined cristae. When investigating temporal gene expression of several mitochondrial-related markers, they indicated fission as the dominant mechanism contributing to these changes observed over time. This is likely to be due to continued mitochondrial stress resulting from the oxidative environment of the retina and prolonged light exposure. We have characterised retinal mitochondrial ageing in a key vertebrate model organism, that provides a basis for future studies of retinal diseases that are linked to mitochondrial dysfunction.

## Introduction

Mitochondria are essential intracellular organelles that provide energy for cells in the form of adenosine triphosphate (ATP), generated through oxidative phosphorylation (Lefevere et al., 2017; Eells, 2019). In addition, they perform a variety of crucial cellular functions including regulating apoptosis, scavenging reactive oxygen species (ROS), calcium homeostasis, nucleotide metabolism, and the biosynthesis of amino acids, cholesterol and phospholipids. Mitochondrial populations are dynamic, showing alterations throughout the lifespan of an organism resulting from fission (division of a mitochondrion into two daughter mitochondria), fusion (merging of two mitochondria into one larger mitochondrion) and mitophagy (selective removal of dysfunctional mitochondria) (Chan, 2020). These processes allow them to maintain cellular homeostasis and meet the energy requirements of the tissue but are also associated with ageing and disease when defective. Examples of retinal disease resulting from defective mitochondria Kearns Sayre syndrome, neuropathy, ataxia and retinitis pigmentosa (NARP), and mito-chondrial encephalomyopathy lactic acidosis and stroke-like episodes (MELAS) (Yu-Wai-Man and Newman, 2017).

Although the diverse functions of mitochondria make them critical for all cell types, mitochondrial populations are most dense in energy intensive tissues like the retina (Eells, 2019; Medrano and Fox, 1995). The photoreceptors consume more oxygen per gram of tissue weight than any other cell type and their inner segment ellipsoids contain numerous mitochondria necessary for high levels of ATP production (Medrano and Fox, 1995; Kam and Jeffery, 2015). Among the mitochondria of the photoreceptor ellipsoid, significantly larger organelles, sometimes referred to as ‘megamitochondria’, have been noted in certain species such as shrews (Lluch et al., 2003; Knabe and Kuhn, 1996) and some teleost species including zebrafish (Nag et al., 2006; Kim et al., 2005). During development, these large mitochondria have been proposed to form from the enlargement of individual mitochondria (Masuda et al., 2016). As the retinal pigment epithelium (RPE) is essential for nourishing the photoreceptors and supporting their high metabolic activity, abundant mitochondria also populate this tissue (Lefevere et al., 2017; Jarrett et al., 2008).

Due to a number of factors, including ROS accumulation caused by elevated ATP generation and light exposure, the mitochondria of the RPE and photoreceptors are particularly vulnerable to dysfunction over time (Upadhyay et al., 2020; Benedetto and Contin, 2019). Alterations in the number and morphology of mitochondria, related to an impaired balance of mitochondrial dynamics, have been observed in ageing retinal tissue and are associated with ocular diseases such as age-related macular degeneration and Leber congenital amaurosis (Lefevere et al., 2017; Jarrett et al., 2008; Toms et al., 2019). Some of these changes can be attributed to mitochondrial fission or fusion. These two processes involve discrete sets of proteins that localise to mitochondria to perform specialised roles. During mitochondrial fusion, Mitofusins Mfn1 and Mfn2 aid the tethering of mitochondria to each other and Opa1 alters the biophysical properties of the lipids in the mitochondrial inner membrane (Formosa and Ryan, 2016). When fission occurs, Fis1 has been proposed to interact with Drp1 to from a collar around the mitochondria that acts to separate them (Liu et al., 2020).

Previously, age-related changes in the retinal mitochondria have been described in several species, including human (Bianchi et al., 2013; Feher et al., 2006), macaque (Gouras et al., 2016), and mouse (Kam and Jeffery, 2015). The zebrafish is a popular model for ocular studies and the ultrastructure of the larval zebrafish photoreceptors and adult retina has been characterised (Tarboush et al., 2012, 2014). However, there has been limited examination of mitochondrial changes throughout the lifespan of the zebrafish retina. In the present study, we have performed ultrastructural analysis to investigate both the development and ageing of mitochondria in the zebrafish RPE and photoreceptors, providing an alternative vertebrate model of retinal ageing.

## Methods

### Zebrafish husbandry

Zebrafish (wild-type, AB strain) were bred and maintained according to local UCL and UK Home Office regulations for the care and use of laboratory animals under the Animals Scientific Procedures Act at the UCL Institute of Ophthalmology animal facility. Zebrafish were raised at 28.5°C on a 14 h light/10 h dark cycle. UCL Animal Welfare and Ethical Review Body approved all procedures for experimental protocols, in addition to the UK Home Office (License no. PPL PC916FDE7). All approved standard protocols followed the guidelines of the Association for Research in Vision and Ophthalmology (ARVO) Statement for the Use of Animals in Ophthalmic and Vision Research Ethics (Westerfield, 2020).

### Mice

Eyes were acquired from mice that had been killed by cervical dislocation in accordance with Home Office (United Kingdom) guidance rules under project license 70/8401. This was done adhering to the ARVO Statement for the Use of Animals in Ophthalmic and Vision Research.

### Transmission electron microscopy

At given timepoints, *z*ebrafish were terminally anaesthetized in 0.2 mg/ml Tricaine (MS-222) and the eyes were harvested through enucleation if 1 mpf or older. Enucleated *z*ebrafish eyes or whole larvae were fixed in 2% paraformaldehyde/2% glutaraldehyde in 0.15 M cacodylate buffer prior to incubation with 1% osmium tetroxide/1% potassium ferrocyanide. Following dehydration in an ethanol series and propylene oxide, the zebrafish were embedded in epon resin. Using a Leica EM UC7 ultramicrotome, 100 nm sections were cut, collected on copper grids (EMS) and stained with lead citrate. Sections were examined on a JEOL 1010 and JEOL 1400Plus TEM, both equipped with a Gatan Orius SC1000B charge-coupled device camera. Images were analyzed using the ImageJ software.

### RT-qPCR

Total RNA was extracted from enucleated zebrafish eyes using the RNeasy micro kit (Qiagen, UK) according to the manufacturer’s instructions. Using 1 μg total RNA, cDNA was reverse transcribed using the Superscript III First-strand synthesis Supermix kit (Thermo Fisher). For quantitative real-time PCR amplifications, gene expression was quantified using SYBR Select fluorescent dye (Thermo Fisher) in triplicate reactions for each sample. All RT-qPCR primers are listed in Table 1. The expression of each gene was normalized to the housekeeping gene *β-actin*. The StepOne Plus RealTime PCR System (Thermo Fisher) was used and reactions analyzed using the Comparative CT experiment option in the StepOne software (Version 2.3).

**Table 1.**
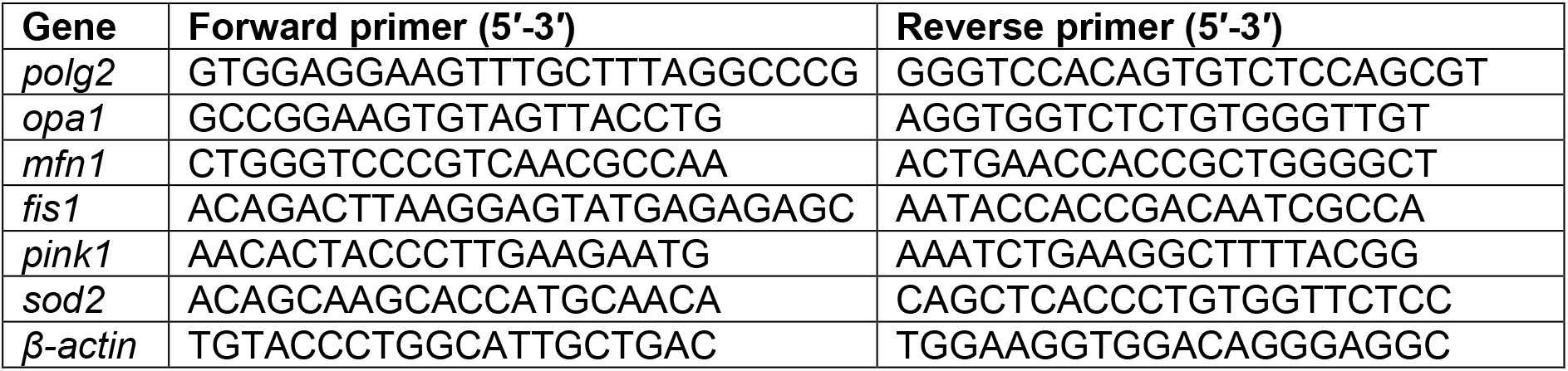
RT-qPCR primers used in this study.

### Statistics

Data are shown as mean values ± standard deviation from n observations. Student’s t-tests were used to compare data. P < 0.05 was accepted to indicate statistical significance (*).

## Results

### Alterations in the mitochondrial morphology within zebrafish retina during development and ageing

#### Retinal pigment epithelium (RPE)

Mitochondria within the RPE were examined in zebrafish from 5 days post fertilisation (dpf) up to 36 months post fertilisation (mpf) by transmission electron microscopy (TEM) (**Figure 1**). In the RPE, a large number of mitochondria were concentrated at the basal surface adjacent to Bruch’s membrane. The RPE of 5 dpf fish contained a small number of notably large mitochondria. By 8 dpf, a dramatic decrease in size and increase in mitochondrial number was observed. With age, the mitochondria continued to decrease in size while increasing in number. Morphologically the mitochondria had a similar appearance at all timepoints except 5 dpf, where they had a more irregular shape and the cristae architecture appeared less well defined.

**Figure 1.**
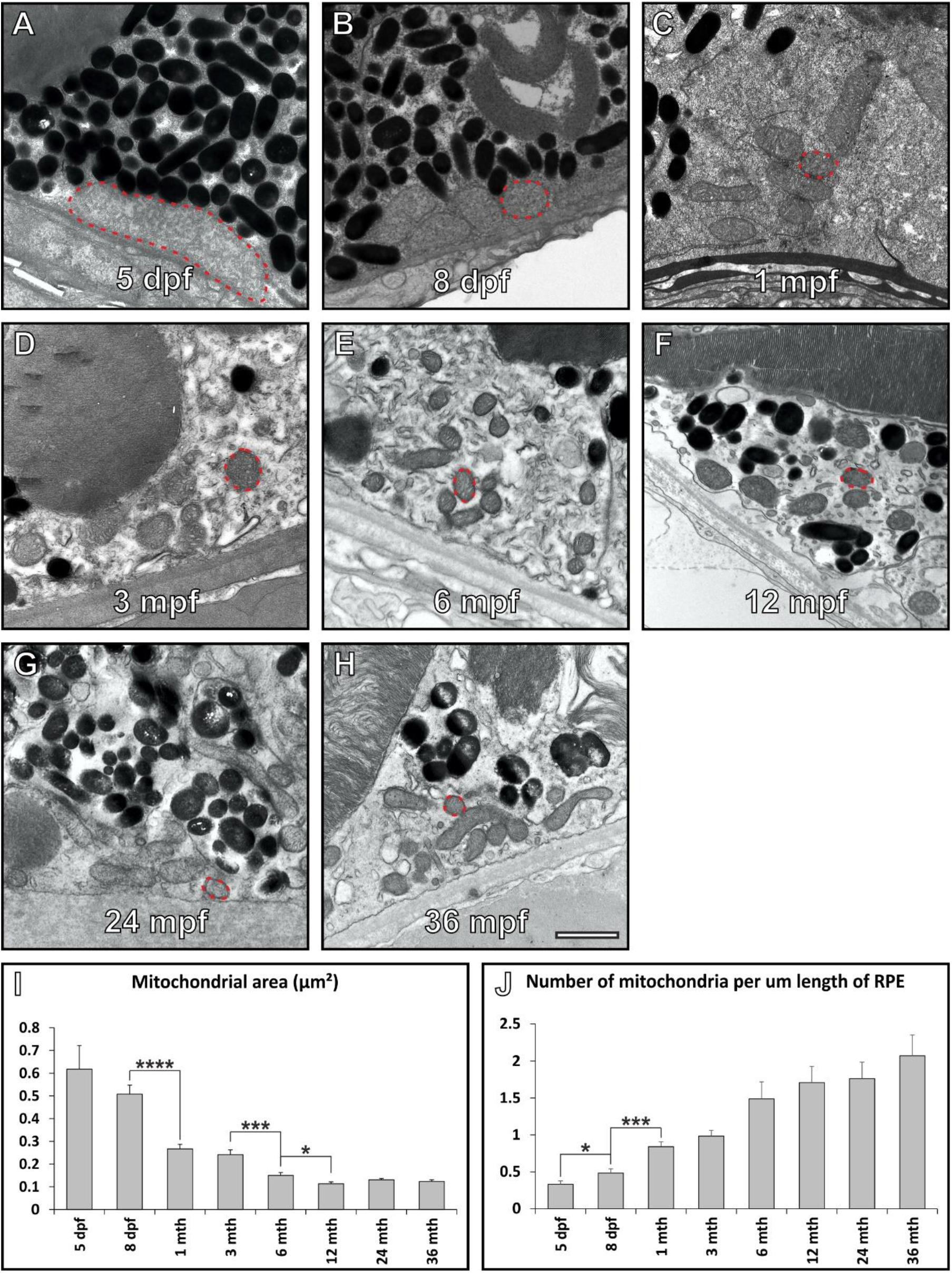
Mitochondria increase in number and decrease in size within the retinal pigment epithelium (RPE) of aging zebrafish retina. (A – H) Electron microscopy images of mitochondria within zebrafish RPE at 5 pdf through to 36 mpf. The red dotted line indicates a mitochondria with the RPE at each timepoint. When examining individual mitochondria within the RPE they (I) decrease in size and (J) increase in number as the zebrafish age. (I – J) At each timepoint N=3 zebrafish were examined. Statistical significance determine by Student’s t-test *P<0.05: ***P < 0.001; ****P < 0.0001. Scale (A – G) 1 μm.

#### Photoreceptor inner segment ellipsoid

Due to difficulties identifying different types of cone photoreceptors and assessing the mitochondria within each, we focused on the morphology of mitochondria within zebrafish rod inner photoreceptors segments (ISs) with age. Mitochondria were examined from 5 dpf up to 36 mpf by TEM (**Figure 2**). At 5 dpf, the rod photoreceptors contained large mitochondria within their ISs bundled together in a spherical arrangement. By 8 dpf, there was an increase in number of mitochondria as well as decrease in size, accompanied by the appearance of more well-defined cristae, reflecting what was observed within the RPE. With age, the morphology of the mitochondria did not appear to alter dramatically.

**Figure 2.**
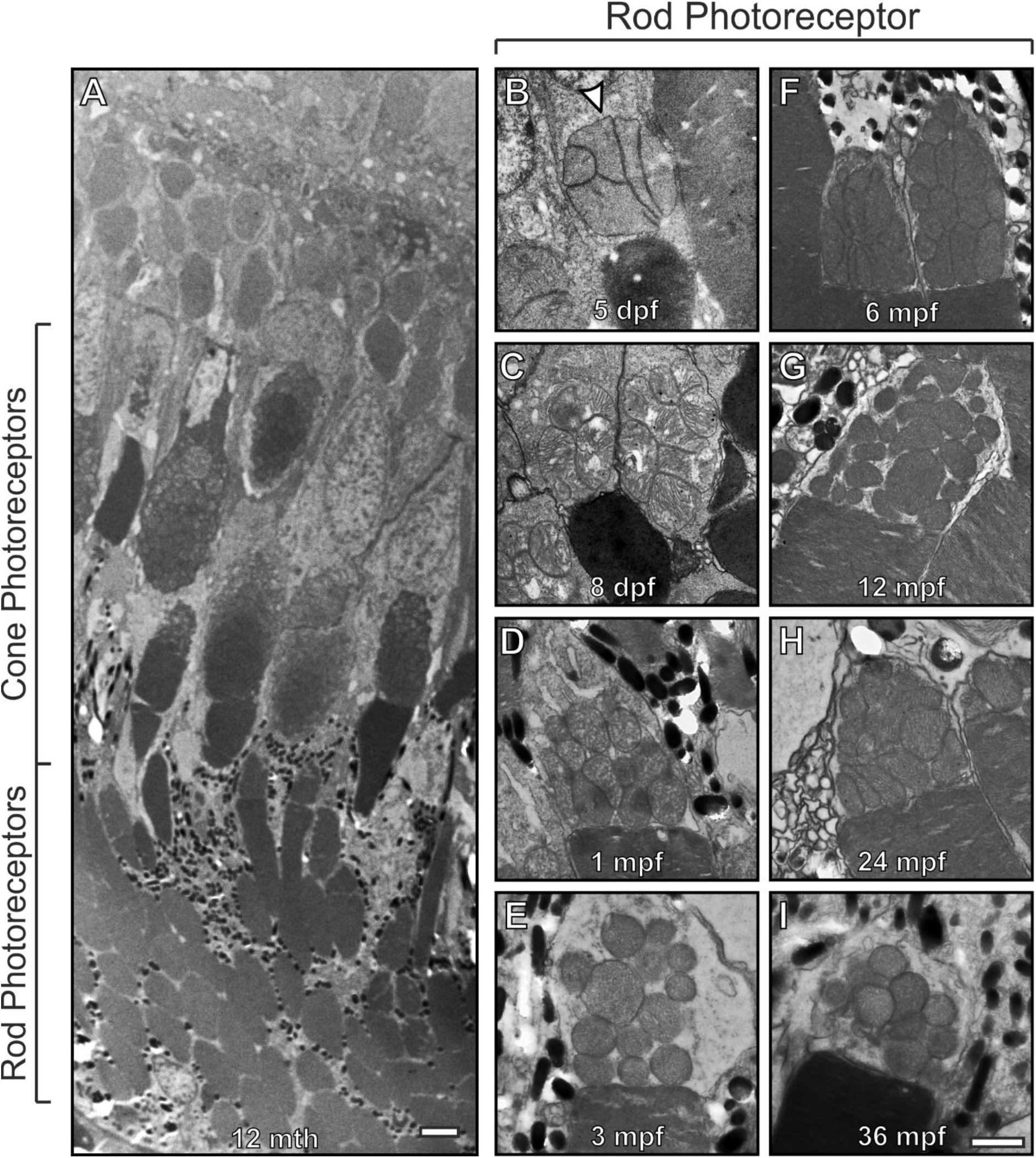
Rod inner segments of early embryonic zebrafish have compact mitochondria with a distant morphology from other ages. (A) Electron microscopy image of 12 mpf retina between the RPE and ONL that includes rod and cone photoreceptor outer (POS) and inner segments. (B – I) Images of rod photoreceptor inner segments at 5 dpf through to 36 mpf. (B) At 5 dpf the mitochondria are bundled together in a spherical like arrangement and there is a morphology change by (C) 8dpf with further changes by (D) 1 mpf as the mitochondria become less clumped together. Scale (A) 10 μm, (B – I) 1 μm.

Cone photoreceptor IS mitochondria were compared at 5 dpf and 1 mpf to determine if a similar change in mitochondrial morphology occurred as observed in the rods (**Figure 3**). At 5 dpf the retina is still developing, and it was not possible to distinguish different cone subtypes unlike at 1 mpf. Even so, the morphology of the mitochondria in 5 dpf cone ISs was clearly different to any of the four cone types identifiable at 1 mpf. At 5 dpf, the mitochondria appeared to be packed together in a spherical like arrangement. Whereas in the fully developed retina at 1 mpf, the mitochondria were still bundled together but more numerous and arranged according to size, with larger mitochondria located more adjacent to the outer segment in the red and green cones, as previously described (Tarboush et al., 2012). At 1 mpf the mitochondria membranes of neighbouring mitochondria appeared closely associated.

**Figure 3.**
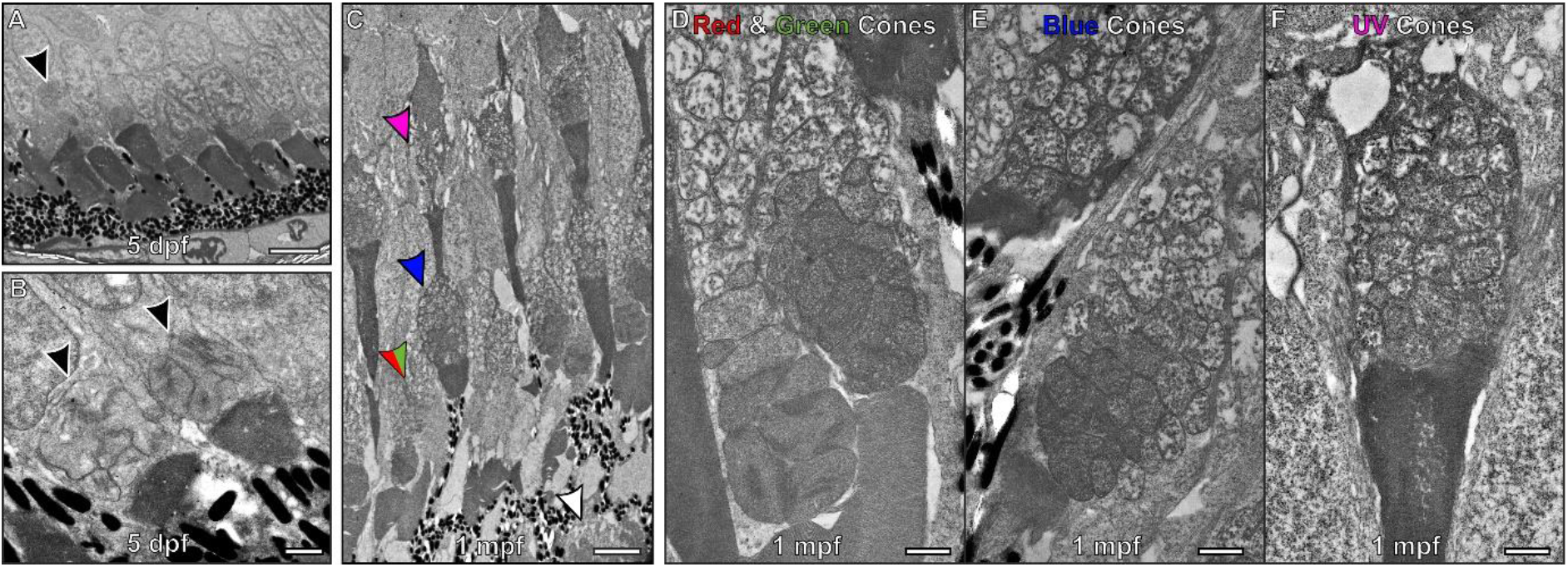
At 5 dpf the mitochondria within cone inner segments have a morphology that is distinct from those found within 1 month old zebrafish. (A) At 5 dpf the retina is still developing and it is difficult to distinguish between cone types. (B) Higher magnification images of cones containing packed mitochondria. (C) By 1 mpf the retina has fully formed and the different photoreceptors can be identified. The arrows highlight the inner segment from different photoreceptors that include red and green arrow for red and green cones, blue arrow for blue cones, purple arrow for the UV cones and the white arrow for rod photoreceptors. (D – F) The corresponding cone types at 1 mpf are shown at higher magnification. Scale (A & C) 5 μm, (B, D-F) 1 μm.

To assess the structure of rod and cone IS mitochondria at 5 dpf in detail, tomograms were generated (**Figure 4 and Video 1 & 2**). These provide higher-resolution data than conventional TEM, in the form of reconstructions that allow mitochondrial membranes to be visualised in 3D. The outer mitochondrial membranes of neighbouring mitochondria were seen to be in close proximity with a uniform spacing between them of 7.66 nm (± 1.43) and 7.67nm (± 1.63) for rods and cones, respectively. Morphological differences between the rod and cone IS mitochondria include more tightly packed cristae in the rods.

**Figure 4.**
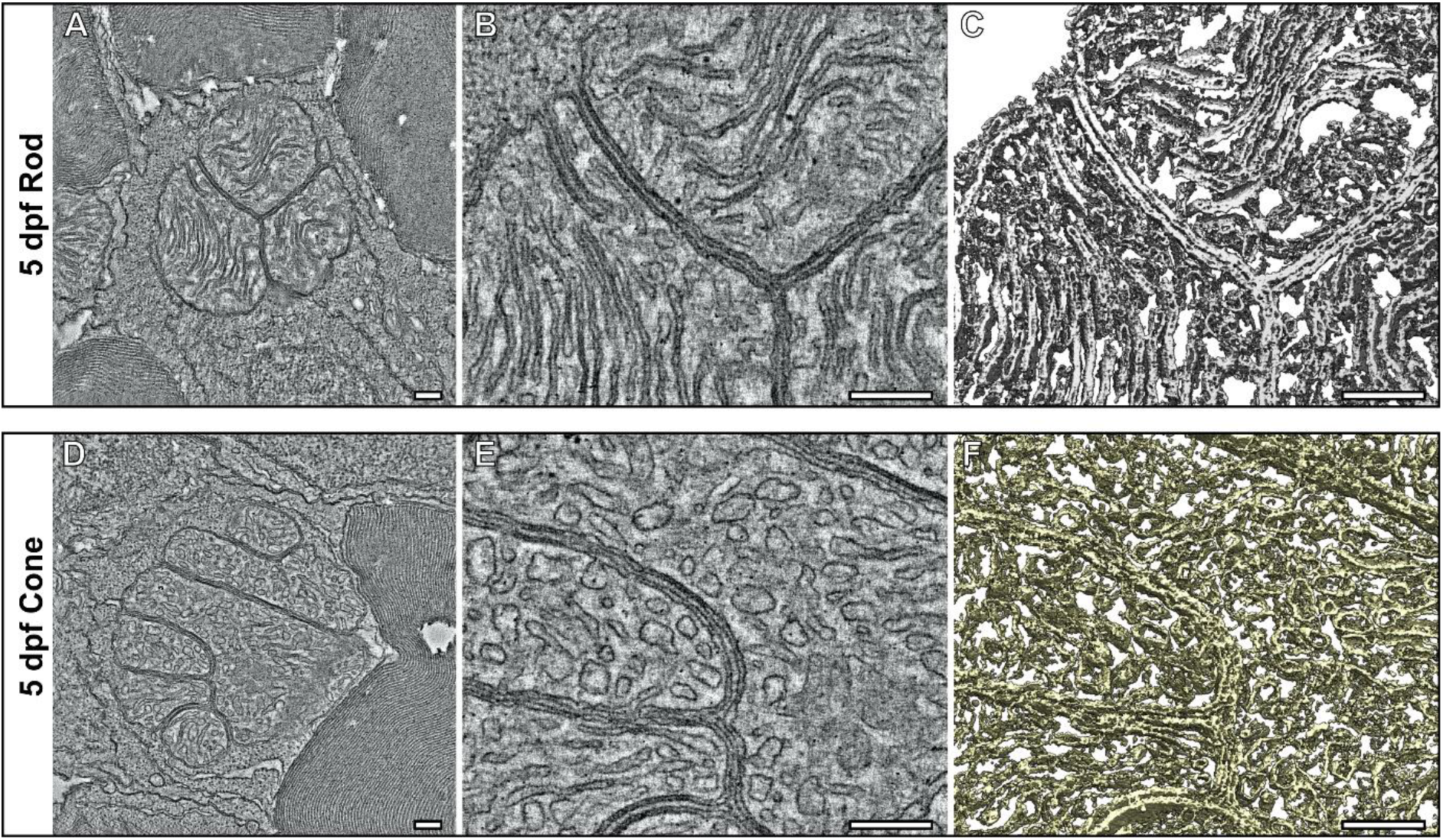
Tomograms shows the compact association of mitochondria within the inner segments of cone and rod photoreceptors at 5 dpf. (A – C) Tomography data of a rod and (D – E) a cone inner segment mitochondria. (A & D) Single slices from the tomograms with (B & E) higher magnification views. (C – F) Surface rendering of the tomography data allow the architecture of the mitochondria to be assessed in 3D. The rod has elongated and tightly associated cristae membranes, where as the cone has cristae that are wider. Scale bars 250nm.

### Morphology of mitochondria within the retina of embryonic and adult mice

To investigate if a notable change in mitochondrial size is seen in the retina of other organisms during retinal development, RPE and photoreceptors of embryonic day 14 (E14), postnatal day 13 (P13) and 6-month-old adult mice were examined (Figure 5). At E14, the photoreceptor outer segments (OSs) had not yet formed and the ISs were positioned up against the RPE layer (Figure 5A - C). As zebrafish develop more rapidly than mice, the rate of mitochondria maturation in the retina most likely differs. The first signs of eye development in mice are detected around E8 and in zebrafish 16 hours post fertilisation (Heavner and Pevny, 2012; Richardson et al., 2017). Therefore, the E14 mouse examined represent an early timepoint in eye development, where retinal mitochondria maturation is still ongoing. P13 mouse retina was more similar in appearance to the 5 dpf zebrafish time-point, as the outer segments are almost fully formed in both (Figure 3A, B and Figure 5D, E). Zebrafish have been reported to have visual function from 3 dpf whereas mice open their eyes at around P11 (Richardson et al., 2017; Rochefort et al., 2009). Therefore, it can be predicted that both 5 dpf zebrafish and P13 mice will have been visually active for ~ 48hrs. When examining the mitochondria within the RPE, there was little difference in their size when comparing the E14, P13 and adult mouse retina (Figure 5 B, E, H). There was a greater difference in the IS mitochondria, at E14 they were shorter in length than at P13 and in adult (Figure 5 C, F, I).

**Figure 5.**
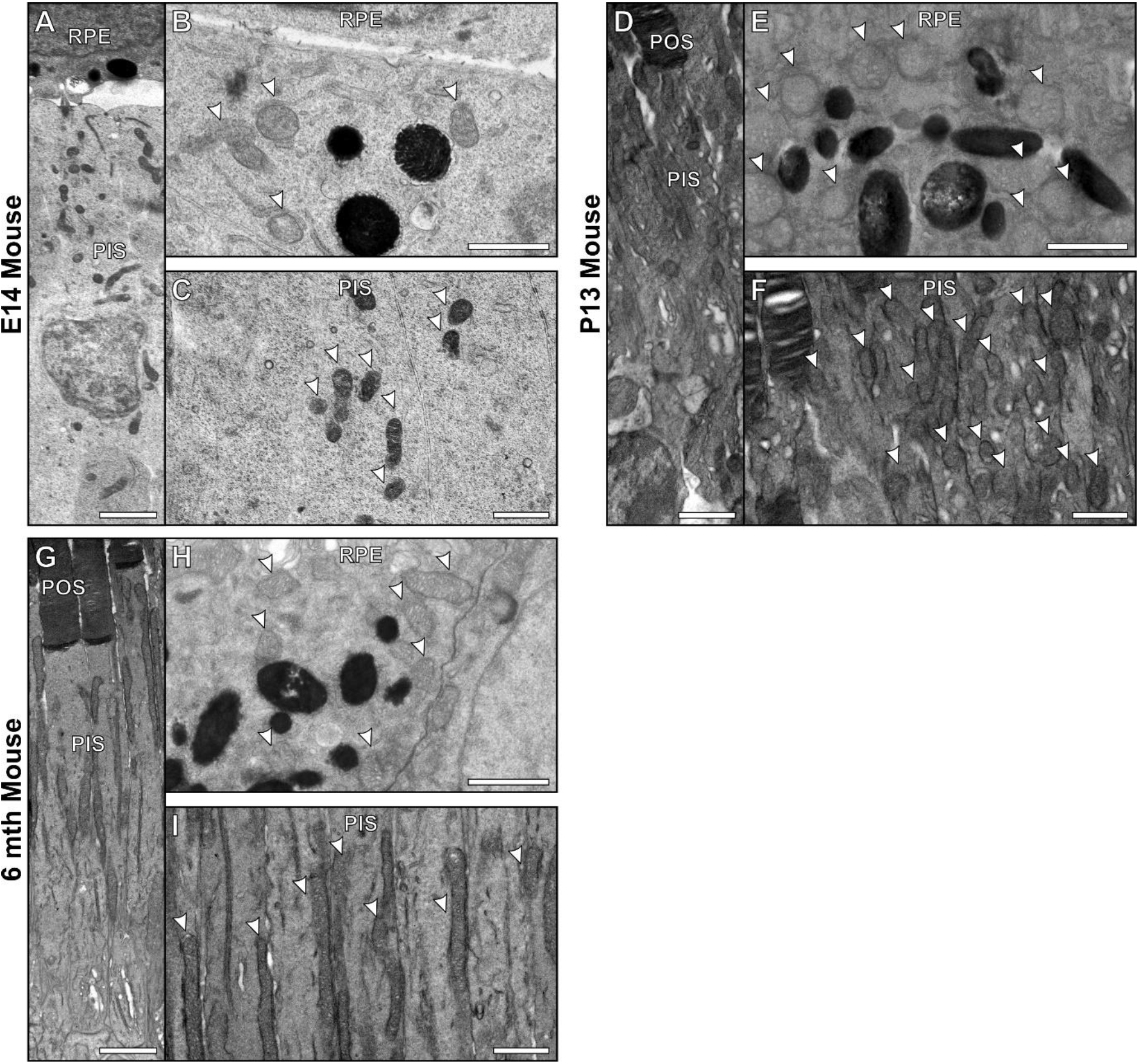
Developing mouse retinas do not have noticeably larger mitochondria within the RPE and photoreceptors compared to adult mice. (A – C) By embryonic day 14 mouse retina is still developing and photoreceptors do not have fully formed outer segments (POS). (D – F) At P13 the mouse have photoreceptors with almost fully developed POS. At P13 the appearance of the RPE and photoreceptors is similar to (G – I) at 6 months in adult mice. There is little difference between the mitochondria within the RPE at (B) E14 compared to a (E) P13 and (H) 6 month old mouse. (C & F) There is greater difference in the photoreceptor inner segment mitochondria. (C) The mitochondria are smaller at E14 compared to (F) P13 and (G) 6 month old mouse. Scale bars (A, D, G) 2μm and (B, C, E, F, H, I) 1μm.

### Age-related changes in mitochondria-associated genes

To examine factors regulating mitochondria size and number in the zebrafish retina, RT-qPCR analysis was performed to assess the gene expression of several indicators of mitochondrial activity: *polg2* (mitochondrial DNA replication), *fis1* (fission), *opa1* (fusion), *mfn1* (fusion), *pink1* (mitophagy) and *sod2* (antioxidant enzyme) (**Figure 6 and 7**). To explore how expression levels change with age, all genes were analysed at 5 dpf, 1 mpf, 12 mpf and 24 mpf. *polg2* showed increased expression at 1 mpf relative to 5 dpf (6.7 ± 1.47 fold; p < 0.01, n = 5) (**Figure 6A**). This increased expression was also present at 12 mpf, (6.4 ±1.26 fold, p < 0.01, n = 5). At 24 mpf, expression of *polg2* was 4.8 ±2.23 fold higher than that at 12 mpf (p < 0.05) (**Figure 6B**).

**Figure 6.**
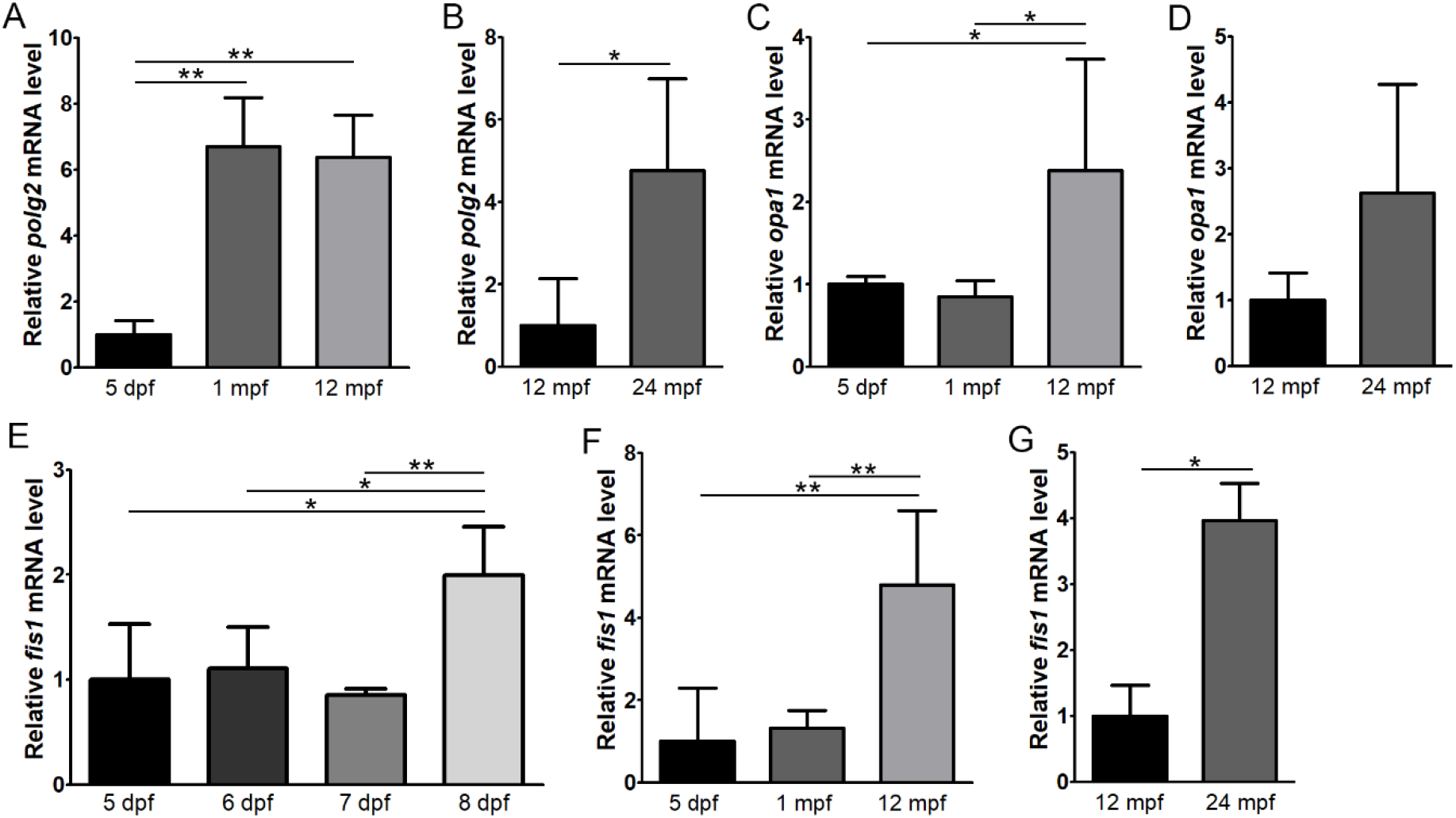
Expression levels of polg2, opa1 and fis1 with age in the zebrafish retina. RT-qPCR was performed to assess expression of polg2 (A, B), opa1 (C, D) and fis1 (E, F) at 5 days post-fertilisation (dpf), 1 month post-fertilisation (mpf), 12 mpf and 24 mpf (n = 5). Expression of fis1 was also assessed at 5 – 8 dpf (n = 4) (G). *p < 0.05, **p < 0.01.

**Figure 7.**
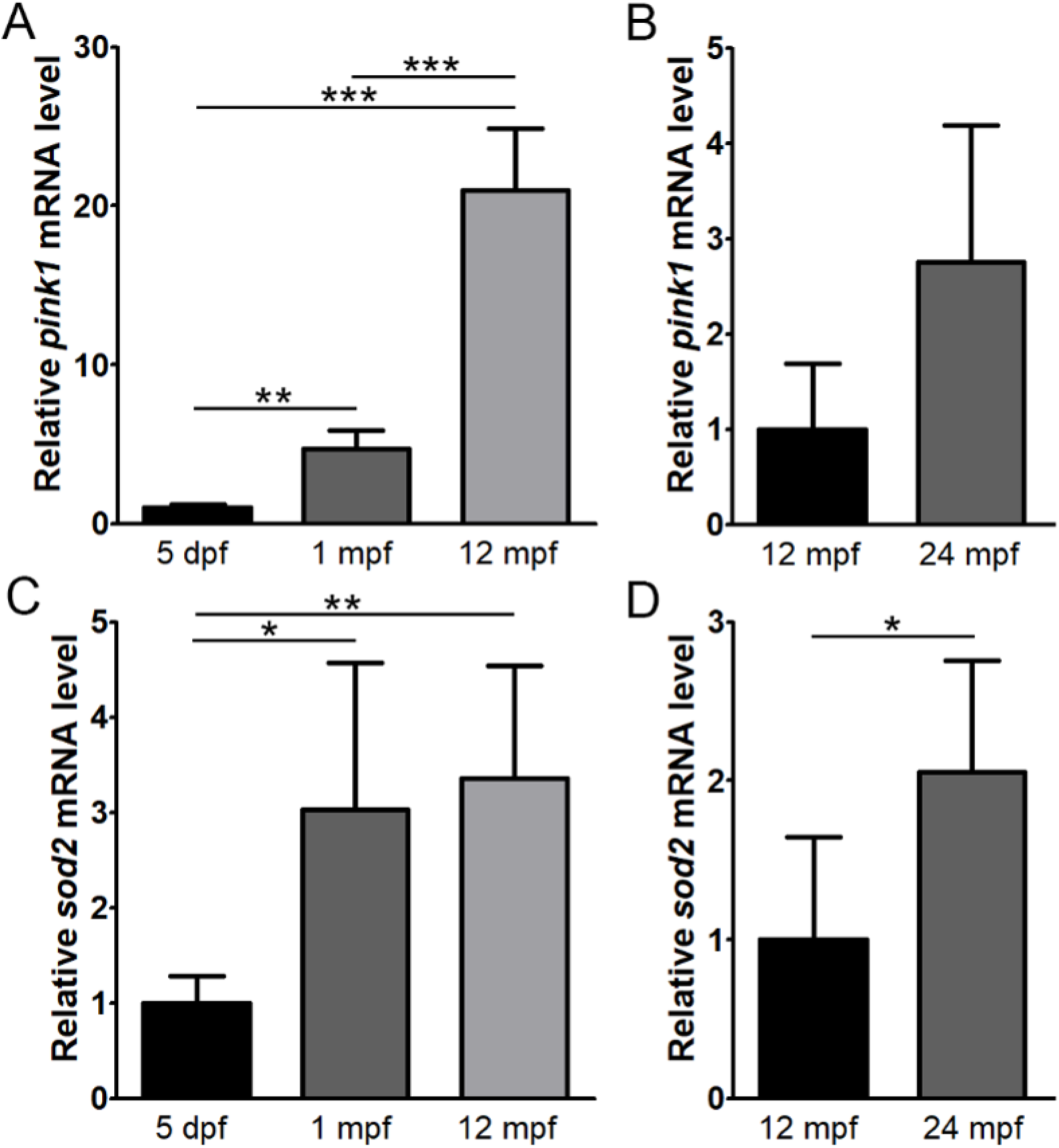
Expression levels of pink1 and sod2 with age in the zebrafish retina. RT-qPCR was performed to assess expression of pink1 (A, B) and sod2 (C, D), fis1 at 5 days post-fertilisation (dpf), 1 month post-fertilisation (mpf), 12 mpf and 24 mpf (n = 5). *p < 0.05, **p < 0.01, ***p < 0.001.

Reduced mitochondrial fusion could lead to the increased mitochondrial numbers and reduced size that was observed with age. However, both markers of fusion, *opa1* and *mfn1*, maintained similar expression levels between the 5 dpf and 1 mpf timepoints (**Figure 6C and Supplementary Figure**). *opa1* expression increased to 2.4 ±1.35 at 12 mpf, which was significantly upregulated compared to both earlier timepoints (p < 0.05). At 24 mpf, *opa1* expression increased, but was not significantly different compared to 12 mpf (p = 0.06). *mfn1* was significantly upregulated at 12 mpf (p < 0.05) and 24 mpf (p < 0.05).

Enhanced mitochondrial fission could also lead to increased mitochondrial numbers and reduced size. At 1 mpf, expression of *fis1* was not significantly different compared to 5 dpf, showing 1.3 ± 0.42 fold relative expression (n = 5) (**Figure 6F**). However, given the substantial increase in mitochondrial numbers that were seen within only 3 days between 5 and 8 dpf (Figure 1), we explored *fis1* expression during this time period. Although it was relatively unchanged at 5-7 dpf, there was a 2.0 ± 0.46 fold increase at 8 dpf relative to 5 dpf (p < 0.05, n = 4) (**Figure 6E**). At 12 mpf, *fis1* expression was upregulated by 4.8 ±1.80 fold compared to 5 dpf (p < 0.01), which was also significantly higher than expression at 1 mpf (p < 0.01) (**Figure 6F**). In addition, expression at 24 mpf (n = 4) was 4.0 ± 0.57 fold greater than 12 mpf (p < 0.05) (**Figure 6G**).

Expression of the mitophagy marker, *pink1*, showed significant upregulation with age, with a 4.7 ± 1.16 fold and 21.0 ± 3.9 fold expression at 1 mpf and 12 mpf, respectively, relative to 5 dpf (p < 0.01 and p < 0.001, n = 5) (**Figure 7A**). The increase in expression between 12 mpf and 24 mpf was not significant (p = 0.09) (**Figure 7B**). *sod2* expression showed a 3.0 ±1.54 fold at 1 mpf compared to 5 dpf (p < 0.05, n = 5) and then maintained a similar level of expression of 12 mpf (**Figure 7C**). At 24 mpf, expression was increased by 2.1 ± 0.71 fold relative to 12 mpf (p < 0.05, n = 5) (**Figure 7D**).

## Discussion

Mitochondria play an essential role in the homeostasis and function of the retina and are intrinsically linked to ageing and disease. In this study, we examined ultrastructural changes in mitochondrial morphology along with associated gene expression patterns to determine how these change in ageing zebrafish retina. This has provided new insights into species specific differences relating to how mitochondria adapt and mature from early development through to later stages of the lifecycle.

As zebrafish age, greater expression of both *opa1* and *fis1* was found within the retina, indicating increasing degrees of mitochondria remodelling with age. Higher relative expression levels of *fis1* were found, compared to *opa1* with ageing, which favours mitochondria fission over fusion. This coincides with an increase in mitochondria number and reduction in their size with age, as measured within the RPE. The most noticeable difference in RPE mitochondria number and size occurred within 1 mpf. At 5 dpf large mitochondria that appear to be precursors becoming smaller and more numerous by 8 dpf. This is likely driven by fission events and coincides with increased expression of *fis1* at 8 dpf that was not seen at 6 or 7 dpf. From 1 mpf through to 36 mpf, the mitochondria continue to reduce in size and increase in number but at a more gradual rate. As fission occurs, mitochondrial DNA is distributed across a larger number of mitochondria and previous studies have indicated this can lead to an increase in mitochondrial DNA replication (Chapman et al., 2020). This may explain the increasing expression levels of *polg2* expressed with age, which encodes p55, an accessory subunit of polymerase gamma involved in mitochondrial DNA replication (Longley et al., 2006). In studies examining the ageing human retina, changes in the size and number of mitochondria within the RPE have been assessed. Between ~40 to 90 years of age, mitochondria were found to reduce in size similar to ageing zebrafish, but in contrast the number of mitochondria reduced with age (Bianchi et al., 2013; Feher et al., 2006).

Zebrafish at 5 dpf had mitochondria with a distinct morphology within the RPE as well as in rod and cone ISs compared to later timepoints. These included large mitochondria within the different cell types and are bundled together with a spherical like arrangement in the photoreceptors. At this timepoint within the RPE and photoreceptors, the mitochondria have an appearance in between the large ones seen at 5 dpf and the smaller mitochondria at 1 mpf. Electron tomography (ET) provided higher resolution data in 3D in comparison to conventional TEM imaging. This allowed detailed assessment of the mitochondrial morphology within photoreceptor ISs at 5 dpf. The arrangement of the cristae in mitochondria in each cell type differed and may relate to the specialised function and environment of the cells. The rods have long sheet-like cristae with membranes that are closely opposed, whereas the cone mitochondrial cristae have a more tubular-like morphology that has greater spacing between membranes. Uniform spacing of the outer mitochondrial membranes was found in both rod and cone ISs and this likely results from tethering complexes that reside in between opposing mitochondrial membranes. These would act to hold the mitochondria together and keep them a specific distance apart similar to tethers found for contact sites between organelles such as the endoplasmic reticulum and lysosomes or mitochondria (Burgoyne et al., 2015).

The term megamitochondria has been used to describe mitochondria that have diameters that exceed 2 μm. This includes the large mitochondria of the photoreceptor in some species of shrew as well as zebrafish (evident in all zebrafish ages examined in this study) (Knabe and Kuhn, 1996; Lluch et al., 2003; Kim et al., 2005). Previous work has shown the megamitochondria in zebrafish photoreceptors generate high levels of energy that are likely to be required to help fulfil the substantial energy demand of these cells (Masuda et al., 2016). Megamitochondria have also be reported to as an indicator of disease as they can occur in hepatic dysfunction (Tandler and Hoppel, 2013). Here we report the existence of physiological megamitochondria within the RPE cells for the first time. It is likely they have not been described previously due to being replaced after 5 dpf with smaller mitochondria. Further work is required to understand why these megamitochondria exist in the RPE, particularly to examine the mechanism that underlies the biogenesis of these organelles.

Changes in mitochondrial morphology with age within zebrafish retina could result from adaptations to compensate for oxidative stress as a result of continued light exposure and the mechanisms that underlie the visual cycle. Light exposure leads to oxygen-related stress of retinal cells including the photoreceptors and RPE as shown with constant LED lighting studies using rats and cell culture models (Godley et al., 2005; Benedetto and Contin, 2019). Phagocytosis of photoreceptor outer segments by the RPE exposures the cells to high levels of free radicals on a daily basis (Upadhyay et al., 2020). Oxidative damage has been shown to cause mitochondria DNA modifications that can lead to mitophagy (Godley et al., 2005; Dan et al., 2020). The expression levels of *pink1* and *sod2* were examined as indicators of mitochondrial stress and an oxidative environment. *pink1* encodes a mitophagy protein required in the selective degradation of damaged mitochondria and *sod2* has been implicated in cardioprotection during oxidative stress by converting superoxide into less damaging species (Jin and Youle, 2012, 1; Flynn and Melov, 2013, 2). *pink1* showed greatly increased expression at 12 mpf indicating continued stress and turnover of mitochondria within the retina that increases with time. Whereas *sod2* had a less substantial increase but still indicated prolonged oxidative stress of the retina compared to the early timepoint at 5 dpf. These observations fit with cellular stress within the retina that increase with age. As individual mitochondria become less efficient due to increasing oxidants with age, upregulated fission could compensate by generating additional mitochondria to fulfil the energy demands.

Developing and mature mouse retinas were examined to determine if a similar morphological change in RPE and photoreceptor mitochondria were evident. Mice mainly have rod photoreceptors and by E14, mouse retinas do not have fully developed photoreceptor OSs. By P13 the mouse retina has a similar appearance to the 5 dpf zebrafish (the earliest point we examined) as the OSs has almost fully formed in both. When examining mouse retina at E14 and P13 and comparing to 6-month adult retina, there was no clear difference in morphology of the RPE mitochondria. In contrast to zebrafish, mouse photoreceptor ISs mitochondria appeared to increase in size between the earliest timepoint examined (E14) and in adult retina (6 months). It is known that mitochondria within photoreceptor ISs change during retinal development, including repositioning of the mitochondria up to postnatal day 21 as shown previously (Meschede et al., 2020). Even so, they do not present the same packing of mitochondria as seen in developing zebrafish photoreceptor ISs. Mice IS mitochondria are orientated along the long axis of the IS with most running alongside and in contact with the plasma membrane (Meschede et al., 2020). Whereas zebrafish IS are more densely packed with mitochondria that are position against the plasma membrane but also throughout the IS and their membranes are closely associated with each other. This dense packing of mitochondria throughout the IS of zebrafish appears to be more similar to human than mouse photoreceptors (Nag and Wadhwa, 2016). Further work is required to determine if the packing of mitochondria in the developing zebrafish photoreceptor ISs and fission into smaller mitochondria in these cells and RPE occurs in other species.

This work provides an overview of the changes in mitochondria morphology and gene expression in developing and ageing zebrafish. The ability for mitochondria to adapt to their surrounding during ageing is essential for cellular health and function. Changes in the mitochondria detected in zebrafish in this study provide an excellent basis for future studies of retinal diseases that are linked to mitochondrial dysfunction.

## Acknowledgments

This research was funded by Wellcome Trust, grant numbers 205174/Z/16/Z (M.M) and 093445 (C.E.F), Sight Research UK, Moorfields Eye Charity (M.M).

