## Supplementary figure for "Fission of megamitochondria into multiple smaller well-defined mitochondria in the ageing zebrafish retina"

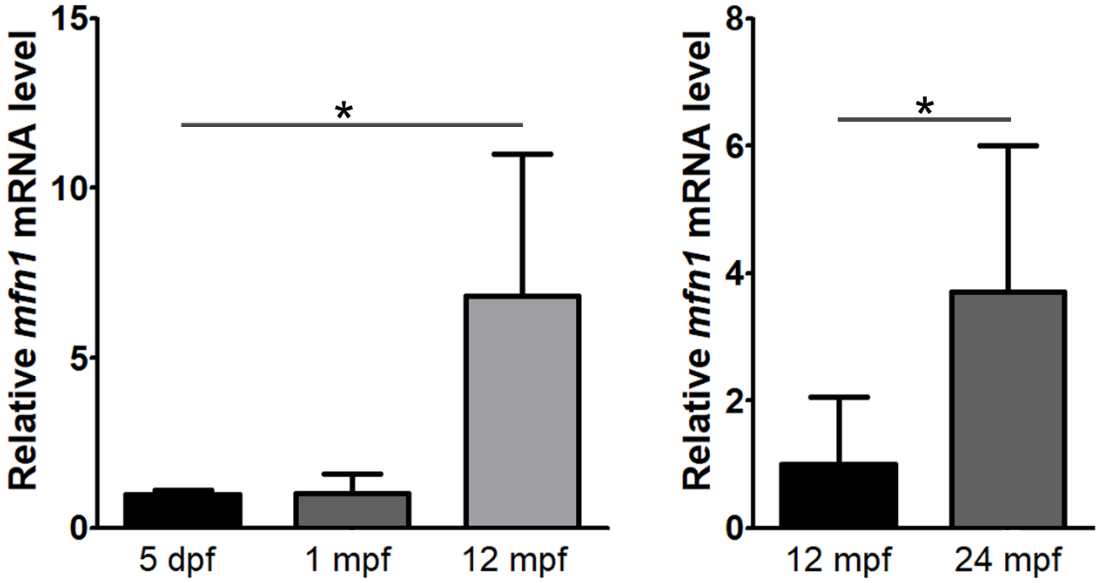


**Expression levels of *mfn1* with age in the zebrafish retina.** RT-qPCR was performed to assess expression of *mfn1* at 5 days post-fertilisation (dpf), 1 month post-fertilisation (mpf), 12 mpf and 24 mpf (n = 5). *$p < 0.05$.
